# High-throughput targeted paleoproteomics sex estimation on medieval Great Moravia individuals using MALDI-CASI-FTICR mass spectrometry

**DOI:** 10.64898/2026.02.17.706309

**Authors:** Fabrice Bray, Anežka Pilmann Kotěrová, Lisa Garbé, Marc Haegelin, Benoît Bertrand, Kevimy Agossa, Christian Rolando, Petr Velemínský, Jaroslav Brůžek, Marine Morvan

## Abstract

The estimation of the biological sex of archeological remains is crucial information in bioarchaeology and forensic anthropology. In recent years, proteomics based on molecular sexual dimorphism have emerged as a preferred method, particularly because of its minimally-invasive approach to extracting amelogenin X and Y proteins from tooth enamel. However, there is an increasing demand to accelerate this process while facilitating the analysis of large archaeological assemblages. This study presents a novel high-throughput targeted paleoproteomics method for biological sex estimation using MALDI-CASI-FTICR mass spectrometry. This approach combines the strengths of existing methods, including ultra-high resolution, significantly reduced processing times, targeted analysis, and scalability to large archaeological sample sets. The method was initially validated on modern individuals with known sex and subsequently applied to 130 adult and juvenile individuals from medieval Great Moravia (present-day Czech Republic). Biological sex was successfully estimated for all but one of the individuals. The results not only provide a more efficient biological sex estimation but also help to resolve a few errors in sex assessment previously encountered with osteomorphological and tooth morphometric techniques. The implementation of this method significantly improves the accuracy and efficiency of biological sex estimation, offering a powerful tool for anthropological research.

**Graphical Abstract:** 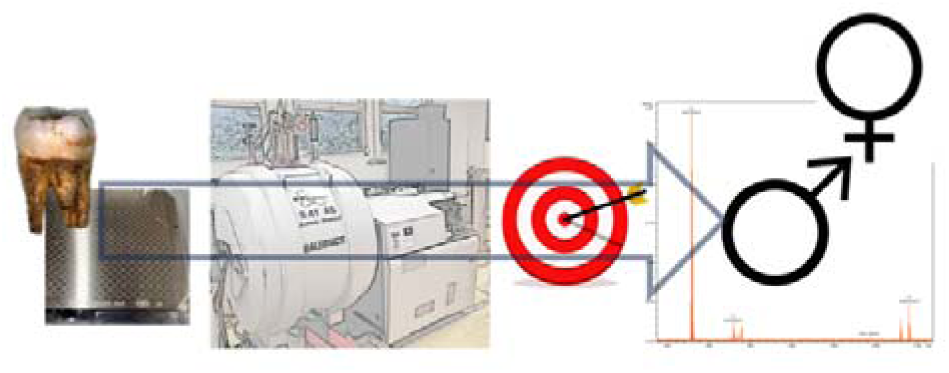

## Introduction

The estimation of the biological sex of human skeletal remains is a core component of both bioarchaeological inference and forensic identification of human remains. Sex estimation provides a structural basis for developing other key elements of the biological profile. Morphological sex assessment is not fully independent, and the validity of morphological assessment is enhanced by the reliable estimation of other biological parameters, particularly age-at-death and ancestry/population affinity. It is well established that the reliability of skeletal sex assessment is sensitive to multiple sources of variation and bias, including inter-and intra-population differences, ontogenetic effects, pathological conditions or taphonomic alterations that may obscure morphological sex assessment. In particular, it is generally inadvisable to attempt morphological sex assessment in fetal, infant, and juvenile remains, as valid and sufficiently accurate osteological methods are not available owing to the limited expression of sexual dimorphism before puberty^1^. In cases where skeletal sex assessment is not feasible or is expected to be unreliable, such as highly fragmentary assemblages or subadult remains, molecular or biochemical methods may provide a useful alternative.

Proteomics approaches based on molecular sexual dimorphism have emerged as powerful tools in recent years^2^, particularly through the minimally-invasive recovery of sex-linked amelogenin (AMELX/AMELY) peptides from tooth enamel^3^, applicable to both adult and subadult individuals^1,4^. Owing to its limited invasiveness and the superior preservation of enamel proteins relative to DNA^2^, proteomic analysis has become a preferred method for biological sex estimation. Amelogenin, a sex-specific protein present in tooth enamel, occurs as two isoforms—AMELX and AMELY—encoded by the X and Y chromosomes, respectively. Using mass spectrometry, peptides derived from these isoforms can be reliably distinguished, enabling accurate estimation of biological sex^5^. This approach has also proven effective in resolving sex attribution errors inherent to osteomorphological assessments^3,6^.

Currently, there is strong pressure to accelerate the entire process of biological sex estimation while simultaneously enabling the analysis of a large number of samples per day^7–11^. Recent Liquid Chromatography-High Resolution Mass Spectrometry (LC-HRMS) studies have demonstrated that modern workflows based on minimally destructive enamel peptide extraction via acid etching combined with targeted proteomics allow for high-throughput (up to 200 samples per day) with high quantitative accuracy and reliable sex estimation, even in archaeological material^7,8^. Another targeted LC-HRMS study showed the possibility of analyzing up to 480 samples per day, but without the use of commercial standards^12^. These methods reduce total analysis time to a few minutes up to tens of minutes per sample without compromising the number of identified peptides or the quality of biological sex estimation. These approaches pave the way for the analysis of entire populations rather than focusing on individual well-preserved specimens.

Both untargeted^3,13,14^ and targeted^7,12,15,16^ approaches have recently been used for biological sex estimation. The untargeted approach allows an exploratory study due to the analysis of the amelogenin sex-dependent peptides, and all other proteins and peptides present in the sample, including contaminations, which, in the case of archaeological samples, can represent a significant ratio. Conversely, the targeted approach allows only preselected amelogenin sex-dependent peptides to be analyzed, regardless of contamination degree. This approach offers the possibility of quantifying peptides of interest for improved biological sex estimation.

The use of commercial standard peptides strategy allows for the most confident sex estimation. Two different approaches have recently been used. The first, external calibration, used by Koenig *et al*.^7^, involves measuring synthetic peptide standards with known concentrations to generate a calibration curve, which is then used to identify the targeted peptides in the analyzed samples. This approach is simple and effective for purified matrices but is sensitive to instrumental variations during the successive analyses of large cohorts. The second method, internal calibration, used by Froment *et al*.^16^, involves measuring synthetic isotope-labeled peptide standards with known concentrations to generate a calibration curve. A known and controlled concentration of isotope-labeled synthetic peptide standards is also added to each sample and can be distinguished by mass spectrometry through their mass shift due to isotopic enrichment. The biological sex estimation was based on the ratio of the intensity of the targeted peptide (light, into the sample) to that of the internal isotope-labeled peptide (heavy, controlled addition into the sample). This approach provides higher accuracy and reliability, can be applied to unpurified matrices, and is not sensitive to instrumental variations; however, the preparation requires an additional step than external calibration.

Matrix-Assisted Laser Desorption/Ionization Mass Spectrometry (MALDI MS), for its part, is rarely used for the estimation of biological sex. Two studies reported the use of Matrix-Assisted Laser Desorption/Ionization-Time-of-Flight (MALDI-TOF)^17,18^ or, more recently, Liquid Atmospheric Pressure-Matrix-Assisted Laser Desorption/Ionization-Time-of-Flight (LAP-MALDI-TOF)^9,10^ for untargeted biological sex estimation of human teeth remains.

Here, we present the first study using a high-throughput targeted Matrix-Assisted Laser Desorption/Ionization coupled with Continuous Accumulation of Selected Ions and Fourier Transform Ion Cyclotron Resonance Mass Spectrometry (MALDI-CASI-FTICR MS) method for biological sex estimation of archaeological remains with internal standards. This method combines the advantages of all methods commonly used to date, including extremely high resolution, significantly shorter processing time, high suitability for highly degraded material, and insensitivity to instrumental variations. These advantages enable the rapid and efficient screening of large sample sets. The targeted MALDI-CASI-FTICR MS workflow further enhances the analytical sensitivity and selectivity, facilitating reliable internal identification of the sex-specific amelogenin peptides AMELX and AMELY. Together, these features establish a rapid, high-throughput, and analytically robust alternative for sex estimation in proteomics, particularly well-suited for large archaeological assemblages and minimally destructive sampling strategies. The method was first designed for modern individuals whose sex is known and was subsequently applied to archaeological teeth from two Early Medieval Great Moravian burial sites in the present-day Czech Republic, comprising both adult and juvenile individuals. In this study, we chose the terminology *biological sex estimation* when relating to biological techniques such as proteomics and aDNA, and *sex assessment* when relating to anthropology.

## Materials and methods

### Samples and osteological analysis

Permanent modern teeth (n = 28; 14 males, 14 females) from individuals of known sex, extracted for therapeutic indications in routine dental practice were obtained for assay development. Use for research was based on written informed consent within an approved ethical framework, with voluntary participation, non-impacting clinical care, and samples handled under confidentiality safeguards. Specimens were provided either via the Forensic Taphonomy and Anatomy Unit or through the Faculty of Odontology (University of Lille, France).

The archaeological samples were obtained from two archaeologically significant Early Medieval Great Moravian burial sites (present-day Czech Republic). First, 30 adult teeth originating from the excavation of the 3^rd^ church at the medieval site of Mikulčice (9^th^–10^th^ centuries) in South Moravia were used in the study. The sex and age-at-death of the adult individuals were previously assessed by Stloukal^19,20^. A revised assessment of sex and age was later published by Zazvonilová *et al*.^21^ using various osteomorphological methods. Sex of some individuals was also assessed by Brůžek and Velemínský in 2017 (unpublished) based on primary and secondary sex diagnosis^22^ (Table 2).

**Table 1.** List of isotope-labeled peptides for AMELX and AMELY (AQUA™ Peptides). R* = isotope-labeled arginine, which induces a shift of +10 Da, and (ox) = oxidation of methionine, which induces a shift of +15.99 Da.

| Peptide sequence | Protein name | Uniprot code-isoform | *m/z* heavy | *m/z* light |
| --- | --- | --- | --- | --- |
| SM(ox)IR*PPY | AMELY | Q99218-1 | 889.4491 | 879.4393 |
| SIR*PPYPSY | AMELX | Q99217-1 | 1089.5621 | 1079.5520 |

**Table 2.** Morphological sex assessment of 30 adult individuals from Mikulčice was performed by Stloukal (1963, 1967), Brůžek and Velemínský (2017), and Zazvonilová et al. (2020), and proteomics sex estimation was performed in the present study.

| Grave number | Age (years) | Sex assessment / Sex estimation |  |
| --- | --- | --- | --- |
|  |  | Osteomorphology | Proteomics |

|  | Stloukal<br>19,20 | Stloukal<br>19,20 | Brůžek,<br>Velemínský<br>2017 -<br>unpublished | Zazvonilová<br>*et al.*<sup>21</sup> | Tooth<br>class | Log<sub>2</sub><br>AMEL<br>X | Log<sub>2</sub><br>AMEL<br>Y | This<br>study |
| --- | --- | --- | --- | --- | --- | --- | --- | --- |
| H362 | 40–50 | M | M | ? | I<sub>2</sub> | 21,13 | 19,01 | M |
| H369 | 40–50 | F | F | ? | <sub>2</sub>I | 20,94 | 0,00 | F |
| H392 | 40–50 | M? | - | ? | M | 23,70 | 0,00 | F |
| H401 | 50–60 | M | ? | ? | P<sub>2</sub> | 22,82 | 20,81 | M |
| H423 | 40–50 | M | M | ? | P<sup>2</sup> | 23,33 | 21,77 | M |
| H425 | 40–59? | M |  | ? | <sub>1</sub>P | 23,94 | 22,19 | M |
| H426 | 40–50 | M | M | ? | C | 23,00 | 21,09 | M |
| H429 | 20–39 | M? | - | ? | M | 23,04 | 21,54 | M |
| H439 | 40–50 | M | - | ? | <sup>2</sup>P | 23,59 | 21,60 | M |
| H483 | 40–50 | M | - | ? | P<sup>2</sup> | 23,80 | 22,08 | M |
| H484 | 50–60 | F | - | ? | <sub>2</sub>P | 23,55 | 0,00 | F |
| H487 | 40–50 | M? | - | ? | C | 23,30 | 21,47 | M |
| H493 | 40–50 | F? | - | ? | M | 23,07 | 21,28 | M |
| H495 | 40–50 | M? | M? | ? | C | 23,61 | 21,67 | M |
| H505 | 20–30 | F | F | ? | <sub>1</sub>P | 22,41 | 0,00 | F |
| H507 | 50–60 | M | - | ? | <sup>3</sup>M | 23,27 | 22,21 | M |
| H512 | adult | F? | - | ? | <sub>2</sub>P | 22,81 | 0,00 | F |
| H551 | 40–59 | M | - | ? | C | 23,07 | 21,17 | M |
| H553 | 20–30 | M | - | ? | C | 23,25 | 21,57 | M |
| H563 | 40–50 | M | M | ? | C | 21,82 | 20,23 | M |
| H565 | 20–30 | M? | - | ? | I<sup>2</sup> | 23,94 | 21,59 | M |
| H588 | adult | F? | - | ? | M | 24,00 | 0,00 | F |
| H589 | 40–59? | M | - | ? | M<sub>3</sub> | 23,10 | 21,69 | M |
| H605a | 14–19 | M | - | -- | <sup>2</sup>I | 23,37 | 21,16 | M |
| H626 | 20–30 | F | - | -- | <sub>2</sub>M | 23,17 | 0,00 | F |
| H628 | 40–50 | M? | - | M | M | 24,15 | 21,98 | M |
| H697 | 50–60 | F | - | ? | <sup>2</sup>M | 23,23 | 0,00 | F |
| H701 | 20–30 | M | M | ? | C | 21,80 | 20,24 | M |
| H1082 | adult | F | F | ? | C | 22,51 | 20,20 | M |
| H1182 | adult | F? | F? | F | C | 23,60 | 0,00 | F |

1 Tooth nomenclature: I = incisor, C = canine, P = premolar, M = molar, upper index = upper 2 jaw, lower index = lower jaw, side of the index indicates the side of the tooth (left or right). A 3 capital letter indicates permanent teeth, and a lowercase letter indicates deciduous teeth. A 4 question mark indicates individuals for whom Zazvonilová *et al.* (2020) stated that sex could 5 not be assessed based on the methods used in the study.

Second, 100 juvenile teeth (both deciduous and permanent) from another Early Medieval Great Moravian necropolis located in Rajhrad (9^th^–10^th^ centuries) were analyzed. Osteobiographical analysis was carried out by Hanáková *et al.*^23^; however, sex assessment was not performed for non-adult individuals. In addition, the sex of 13 individuals was previously assessed by Pospíšilová^24^ using the permanent teeth metric method^25^ (Table 3).

**Table 3.**
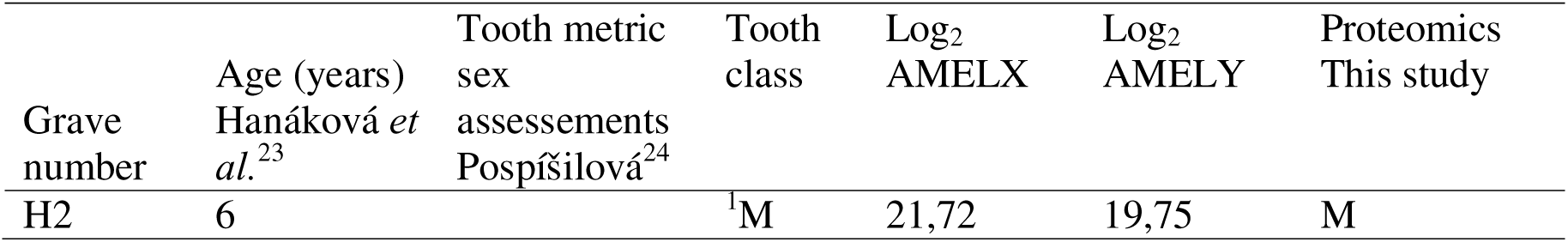
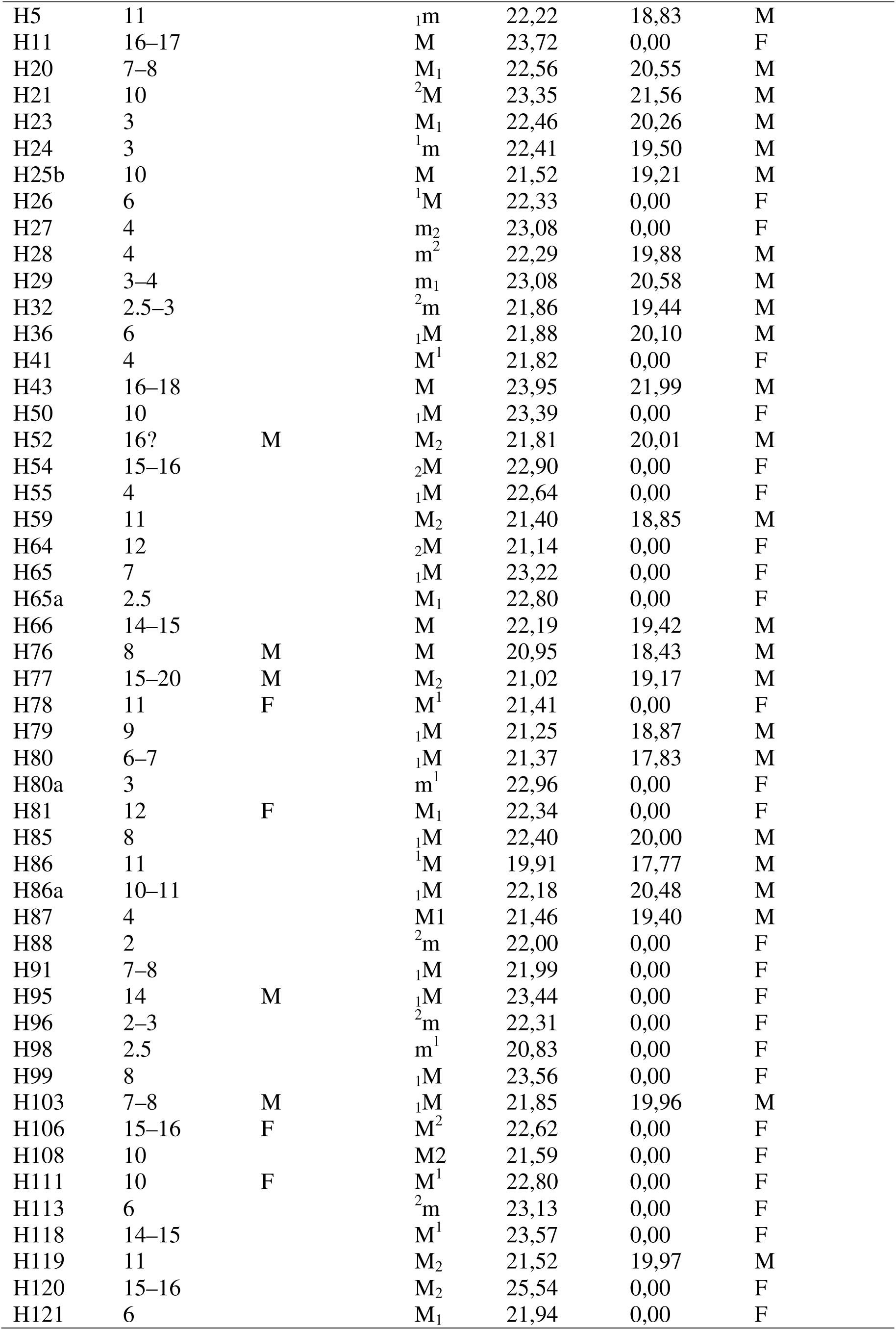
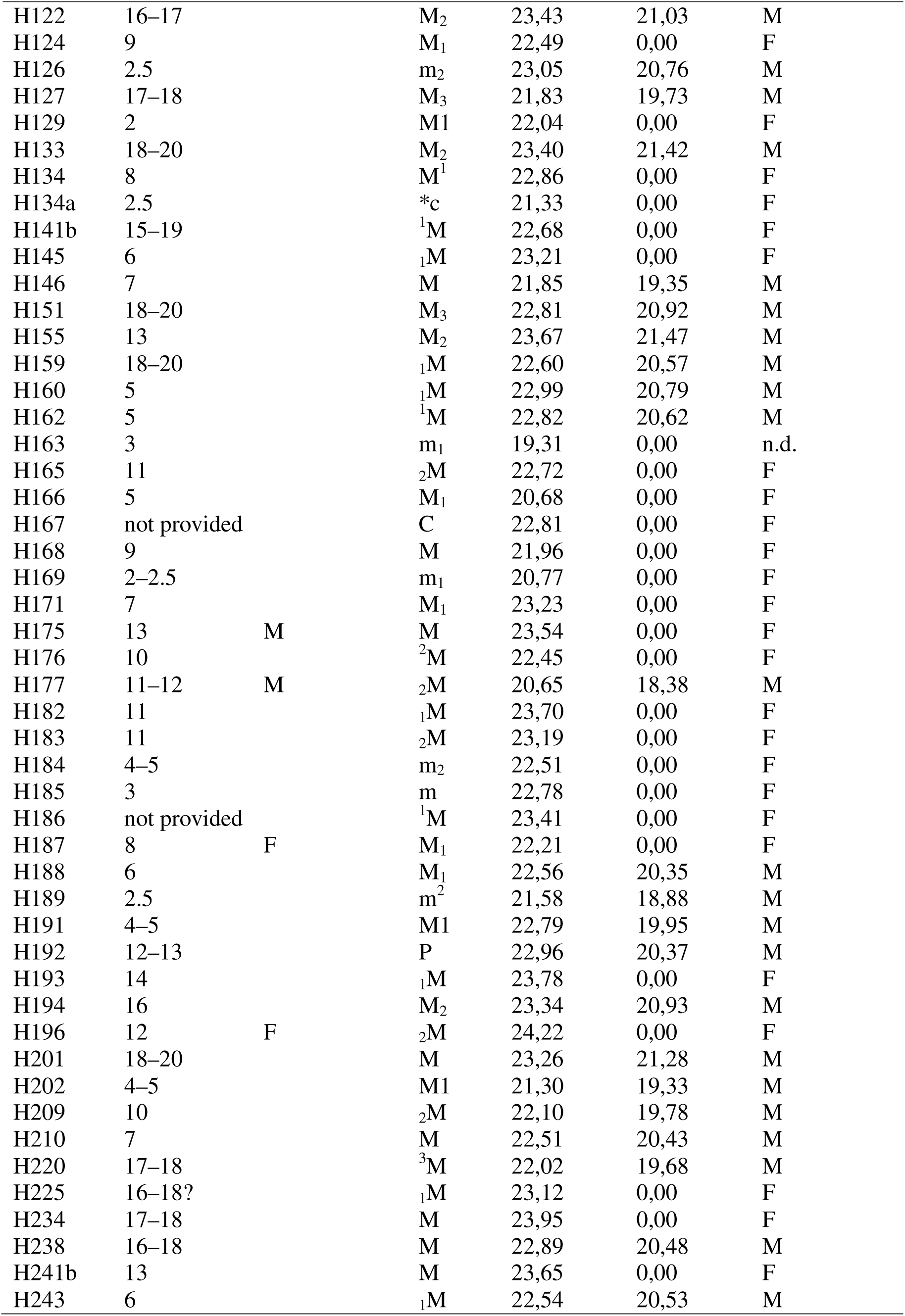
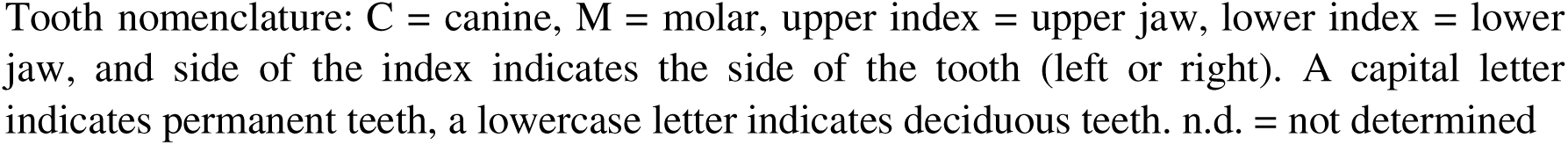
Metric sex assessments of 100 juvenile individuals from Rajhrad based on permanent teeth by Pospíšilová (2017) and proteomics sex estimation performed in the present study.

### Reagents and chemicals

All aqueous solutions were prepared using ultrapure grade water obtained by water filtration with a Direct-Q3 system (Merck Millipore, Burlington, Massachusetts, United States). All chemicals, biochemicals, and solvents were purchased from Merck (Merck KGaA, Darmstadt, Germany) and used without additional purification. All solvents were of MS analytical grade (ThermoFisher, Kandel, Germany).

### Peptide extraction by acid etching and purification

Each tooth was individually cleaned with LC-MS-grade water. The tooth was then incubated in 200 μL of 0.6 N HCl at room temperature for 1 h. The crown of the tooth was incubated in the cap of 1.5-mL microcentrifuge tube. Enamel peptide solutions were then purified using 96-well plates C18 (Affinisep, Petit-Couronne, France). The peptides were resuspended in 10 µL of 0.1% formic acid in H_2_O.

### FTICR MS analysis

The desalted peptides were diluted 2 times in 0.1% formic acid in H_2_O. Diluted peptides (0.6 µL) were deposited on 384 Ground steel MALDI plates (Bruker Daltonics, Bremen, Germany) with 0.6 µL of isotope-labeled peptides (AQUA™ Peptides, Merck KGaA) at 6 fmol for AMELX and AMELY (Table1) and 1 µL of HCCA matrix at 10 mg/mL in ACN/H_2_O (80/20, *v/v*). TFA (0.1%) was added to each sample spot and dried at ambient temperature.

MALDI-FTICR MS experiments were performed on a Bruker 9.4 Tesla SolariX XR FTICR mass spectrometer controlled by FTMSControl software and equipped with a CombiSource and a ParaCell (Bruker Daltonics, Bremen, Germany). A Bruker Smartbeam-II Laser System was used for irradiation at a frequency of 1,000 Hz and using the “Minimum” predefined shot pattern. MALDI-FTICR spectra were generated from 500 laser shots in the *m/z* range from 600.01 to 5,000 with 1 M data points (*i.e.*, transient length of 2.2020 s). Twenty spectra were averaged. The transfer time of the ICR cell was set to 1.2 ms and the quadrupole mass filter operating in RF-only mode was set at *m/z* 600.

MALDI-CASI-FTICR MS spectra were generated using parameters similar to those of MALDI-FTICR MS with some modifications. Fifteen spectra were averaged. Five ions were selected: *m/z* 829.4566 (SIRPPYP, AMELX protein, named AMELX1), 879.4393 (SM(ox)IRPPY, AMELY protein, named AMELYL), 889.4491 (SM(ox)IR*PPY, AMELY protein, named AMELYH), 1079.5520 (SIRPPYPSY, AMELX protein, named AMELX2L), 1089.5621 (SIR*PPYPSY, AMELX protein, named AMELX2H), with isolation windows at *m/z* 10. The acquisition time for each sample was less than 1 min.

### Peptides identification

MS raw data from MALDI-FTICR were processed using DataAnalysis 5.0 software. The FTMS algorithm was employed with the following parameters: S/N > 4, Relative intensity threshold 0.01%, and Absolute intensity threshold 100. The list of peptide markers was searched using a homemade Python program (version 3.12.7, available on GitHub https://github.com/source-code-guru/SearchPPM). The list of [M+H]^+^ masses is indicated in the program, and the search was carried out with an error of 1 ppm. Log_2_ values for AMELX and AMELY intensities and heavy/light peptide ratios were calculated. Samples with Log_2_ AMELX < 20, Log_2_ AMELY < 16, ratio 889/879 > 10, and ratio 1089/1079 > 120 were considered indeterminate. The graphical representations are created using R studio (version 2025.09.2+418).

## Results

### Development of the high-throughput MALDI-CASI-FTICR MS method on modern individuals

Sex estimation by mass spectrometry involves detecting specific peptides of amelogenin isoforms X and Y. Our methods focused on the detection of three peptides (AMELX1, AMELYL, and AMELX2L) derived from AMELX and AMELY proteins used by Adair *et al.*^9^ and two peptides (AMELYH and AMELX2H) used by Koenig *et al.*^7^ were selected to be added to the sample as standards. With the MALDI-CASI-FTICR MS targeted method, peptide masses searched are single-charged, and the five peptides of interest, including AQUA™ Peptides are selected and detected (Figure 1). As shown in Figure 1, the female sample contains the masses *m/z* 829.4553 (AMELX), *m/z* 879.4407 (heavy peptide AMELYH), *m/z* 1079.5520 (AMELX2L), and *m/z* 1089.5621 (heavy peptide AMELX2H). For the male sample, the spectrum contains the peptide at *m/z* 879.4407 (heavy peptide AMELYH).

**Figure 1:**
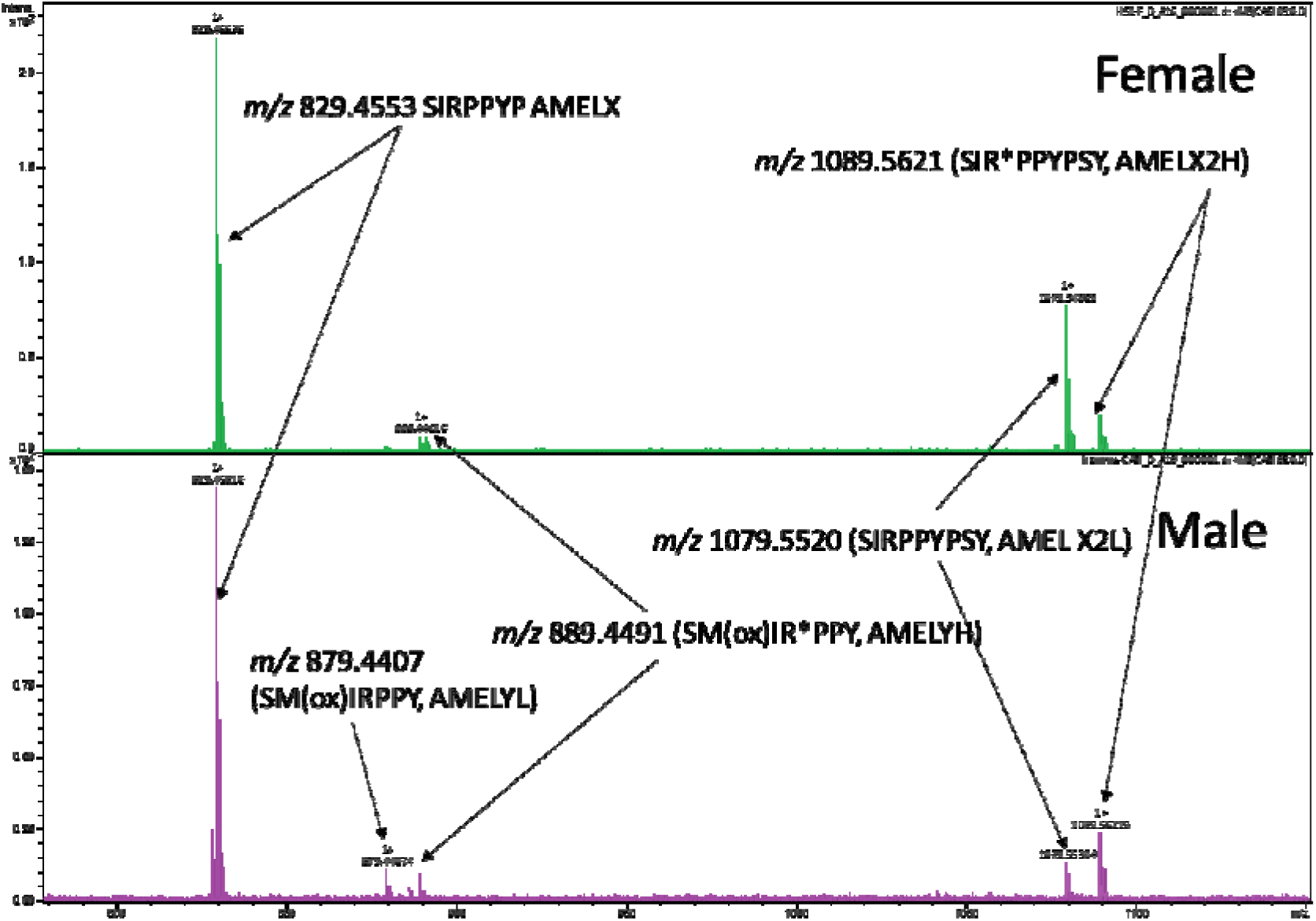
MALDI-CASI-FTICR mass spectra of modern male and female individuals. The peptides AMELX1, AMELX2, and AMELY, as well as the two AQUA™ Peptides, are indicated. Heavy peptides are annotated with H and light peptides with L. The indicated peptides were monocharged.

Supporting Information S1, Figure S1 shows the MS spectra of modern male references obtained with MALDI-CASI-FTICR and MALDI-FTICR MS. In the MS spectrum, numerous masses corresponding to peptides were observed. These masses correspond to peptides from amelogenin X and Y, enamelin, and collagen proteins (Supporting Information S1, Figure S1 A). Zooming in on the region containing the peptides of interest shows that the CASI method allows only the selected peptides to be detected and facilitates their identification (Supporting Information S1, Figure S1 B). With this targeted method, data analysis allows for simpler and faster identification of amelogenin peptides of interest.

A dilution series of AQUA™ Peptides in a pool of five modern male extraction solutions ranging from 0.3 to 600 fmol was analyzed using the MALDI-CASI-FTICR MS method and showed good linearity (R² = 0.989 for AMELY and R² = 0.997 for AMELX) (Figure 2 A, B). To perform quantitative analysis, it is important to determine the limit of detection (LOD) or sensitivity of the method, and the limit of quantification (LOQ) of each targeted peptide. The limit of detection signifies the lowest signal that can be detected, whereas the limit of quantification denotes the lowest MS signal that can be reliably quantified. The LOD for each peptide was 0.6 fmol, and the LOQ was 3 fmol. For the analysis of archaeological samples, each AQUA™ Peptides was spiked at 6 fmol to enable effective detection without affecting the detection of native peptides. Regarding the data on LOD and LOQ, our results are similar to those obtained by LC-HRMS and published by Koenig *et al.*^7^.

**Figure 2:**
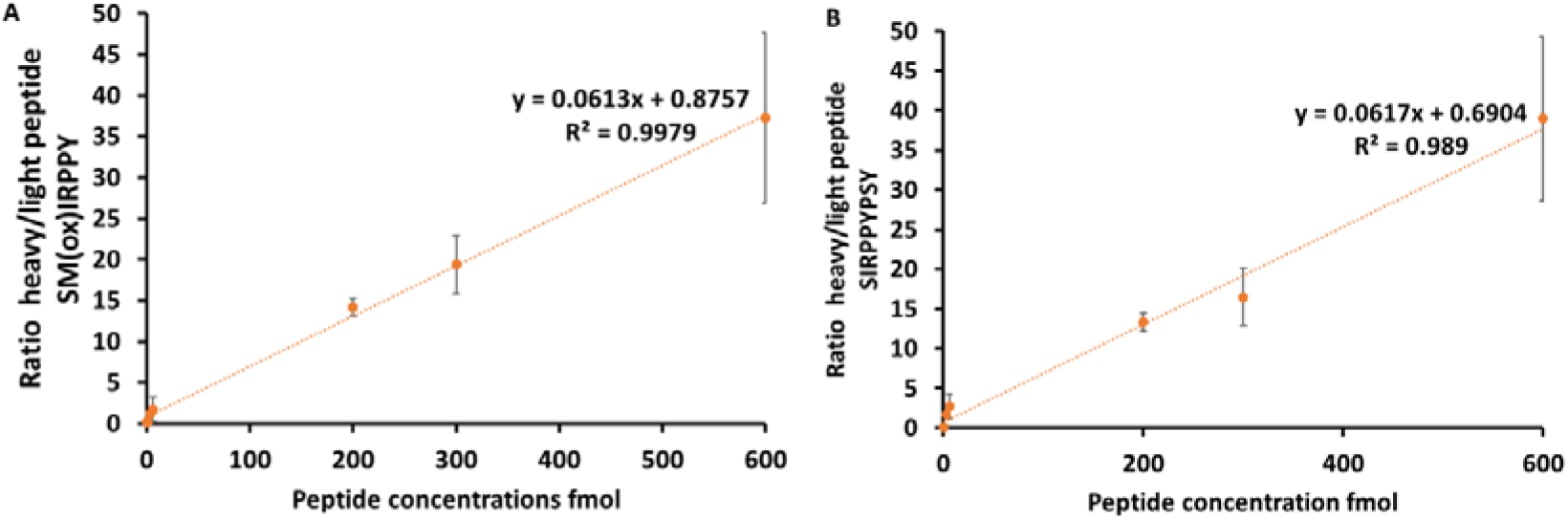
Dilution series of SM(ox)IRPPY, AMELYH (A), and SIRPPYPSY, AMELX2H (B) peptides ranging from 0.3 to 600 fmol in a pool of five modern male extraction solutions using a MALDI-CASI-FTICR MS targeted method. All analyses were performed in triplicate. For each linear regression, the equation and R² are indicated. Black bars correspond to the standard deviations.

Using our modern male reference samples, we observed a linear correlation between Log_2_ AMELX2L (*m/z* 1079.5520) and Log_2_ AMELY (*m/z* 879.4393), with a Pearson correlation of 0.641 (p-value = 0.001). We observed a linear correlation between Log_2_ AMELX1 (*m/z* 829.4566) and Log_2_ AMELY (*m/z* 879.4393), with a Pearson correlation of 0.731 (p-value = 0.0001) (Figure 3).

**Figure 3:**
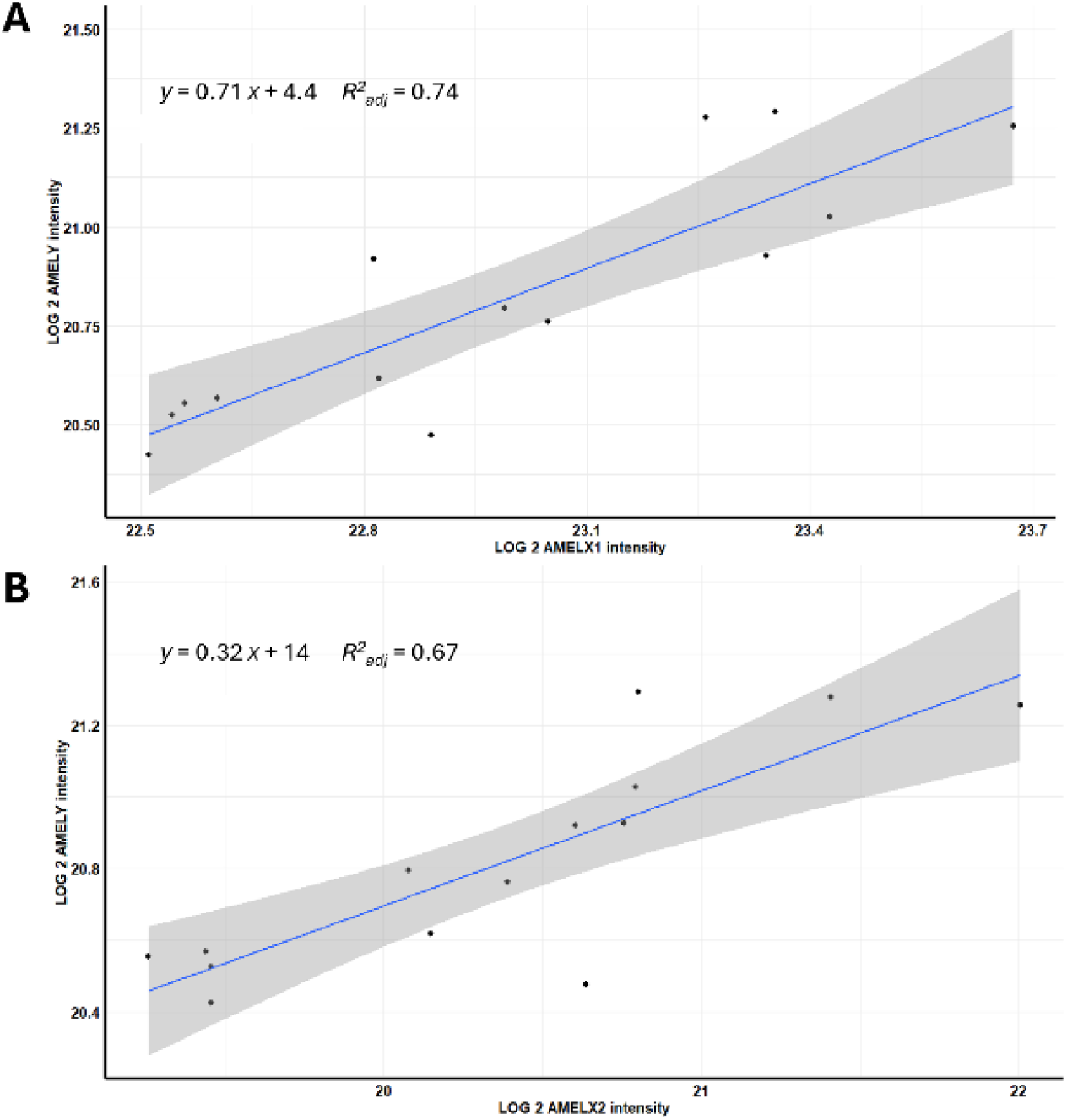
Linear regression of Log_2_ AMELY intensity as a function of Log_2_ AMELX1 intensity (A) or AMELX2L (B). This study involved the analysis of 14 modern males using MALDI-CASI-FTICR. Blue line = linear regression model; grey line = confidence interval (observed values 95%); black dot = sample.

MALDI-CASI-FTICR MS methods allowed the estimation of the biological sex of modern individuals based on Log_2_ AMELY intensity and Log_2_ AMELX1 or Log_2_ AMELX2 intensities (Figure 4). With this method, we have 100% specificity in the cohort of 14 males and 14 females.

**Figure 4:**
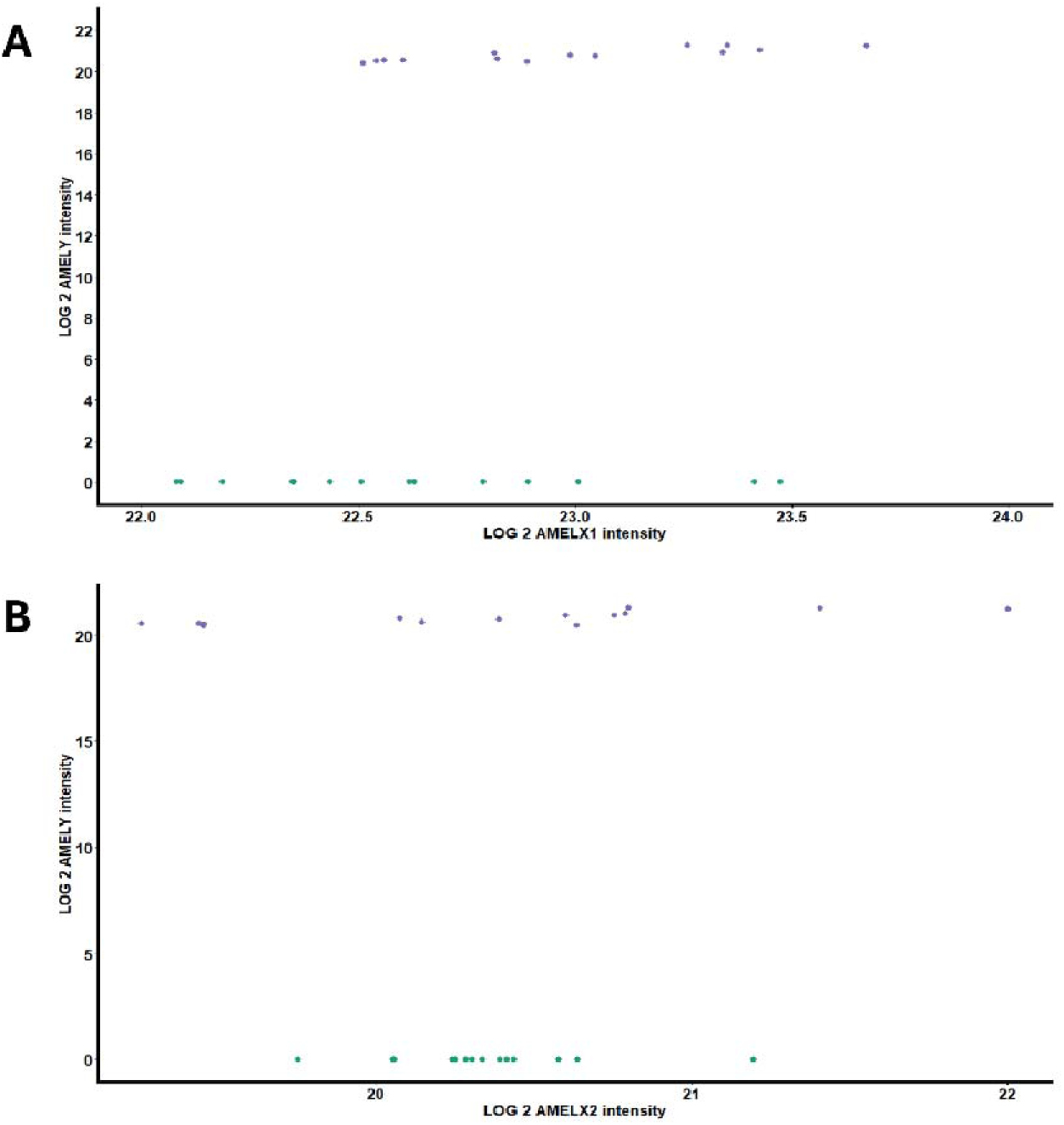
Overview of sex estimation of the 28 modern teeth by MALDI-CASI-FTICR, represented as the Log_2_ AMELY intensity as a function of Log_2_ AMELX1 intensity (A) or AMELX2L (B) for each sample. The blue and green dots correspond to males and females, respectively.

The Log_2_ ratio of intensities for the peptide *m/z* 889.4491 (SM(ox)IR*PPY, AMELY protein, named AMELYH) / *m/z* 879.4393 (SM(ox)IRPPY, AMELY protein, named AMELYL), and *m/z* 1079.5520 (SIRPPYPSY, AMELX protein, named AMELX2L) / *m/z* 1089.5621 (SIR*PPYPSY, AMELX protein, named AMELX2H) were calculated and plotted on a graph (Supporting Information S1, Figure S2).

The results showed a better correlation with the intensities of AMELY and AMELX1. On this basis, threshold values were determined: Log_2_ AMELX < 20 and Log_2_ AMELY < 17. Considering the ratio of light to heavy peptides, if the ratio 889/879 > 10, the sample was considered indeterminate. The 1089/1079 ratio showed a wide variation in intensity. Samples with a 1089/1079 ratio > 120 were considered indeterminate. With our method, heavy peptides were used as internal standard and enabled the detection threshold to be determined in order to avoid false negatives (to estimate a female when it is in fact a male).

### Biological sex estimation of archaeological adult and juvenile teeth

Both high-throughput untargeted and targeted quantitative proteomics MALDI-FTICR MS and MALDI-CASI-FTICR methods were performed on 130 remaining samples from medieval Mikulčice and Rajhrad in Great Moravia. Based on the heavy/light peptide ratio, both methods led to the identification of 60 females, 69 males, and 1 indeterminate individual (Figure 5). The heavy/light peptide ratios for all samples are shown in Supporting Information S1, Figure S3, and Supporting Information S2, Table S1, S2.

**Figure 5:**
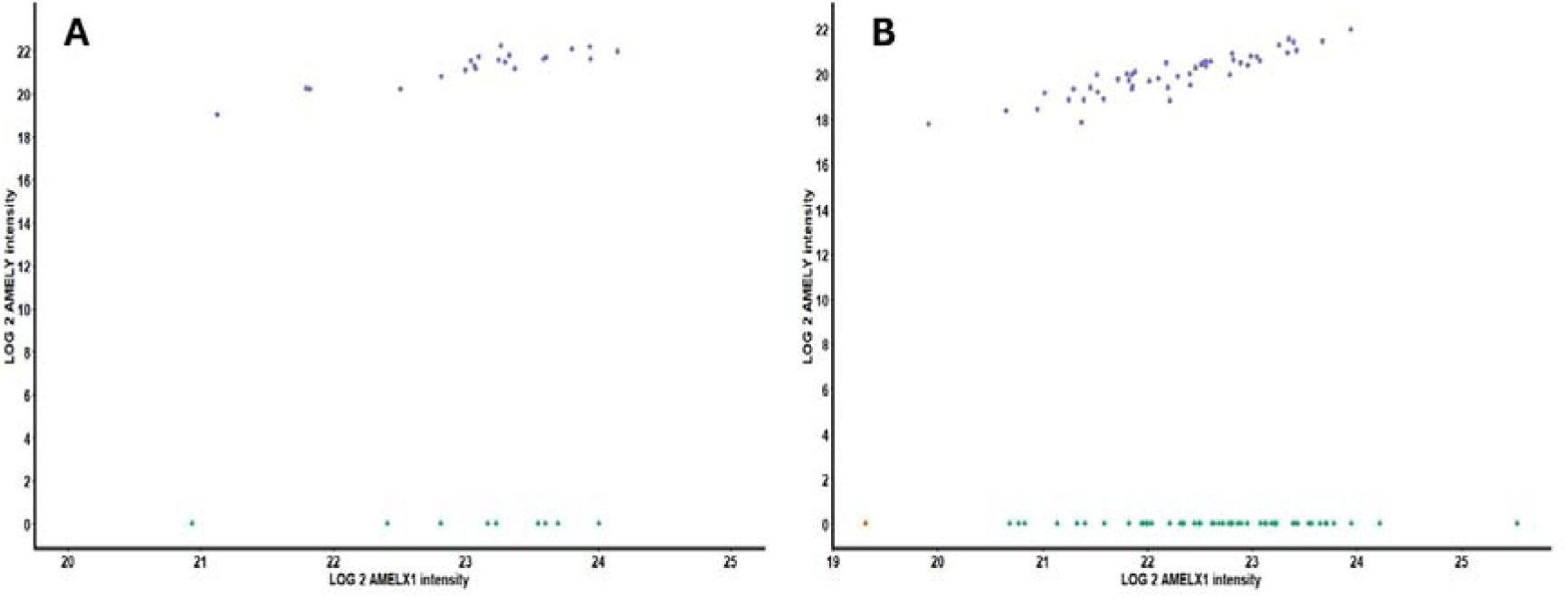
Overview of biological sex estimation for archaeological samples by MALDI-CASI-FTICR, represented as the Log_2_ AMELY intensity as a function of Log_2_ AMELX1 intensity. Biological sex estinmation of the Mikulčice (A) and Rajhrad (B) sites. Blue dots correspond to males, green dots to females, and orange dots correspond to indeterminate samples.

### Comparison of proteomics sex estimation with sex assessments obtained by conventional methods

The sex of 30 adult individuals from Mikulčice was originally assessed by Stloukal^19,20^ based on their skeletal morphological traits. Proteomics sex estimation was consistent with these assessments, except for three individuals (H392, H493, and H1082), for whom the proteomics results indicated the opposite sex (Table 2).

In the case of non-adult individuals from Rajhrad—whose sex is generally not assessed using traditional osteological methods due to the absence of fully developed sexually dimorphic skeletal features—Hanáková *et al.*^23^ did not perform sex assessment. However, Pospíšilová^24^ assessed the sex of non-adult individuals from Rajhrad using a metric analysis of permanent teeth. Among the 13 individuals shared between her study and the present work, we proteomically confirmed the sex of 11 individuals (Table 3).

## Discussion

The MALDI-FTICR mass spectrometer used in this study has the highest resolution compared to other mass spectrometers currently used for biological sex estimation and is becoming the technique of choice. Its MALDI source has the advantage of not using chromatography, which drastically reduces the analysis time and estimated costs. The combination of high-throughput and targeted approaches can confidently estimate the biological sex of up to 1440 individuals per day and represents the most efficient method to date. The use of synthetic isotope-labeled peptide (AQUA™ Peptides) standards allowing for greater consistency and reproducibility between analyses and laboratories (Table 4).

**Table 4.**
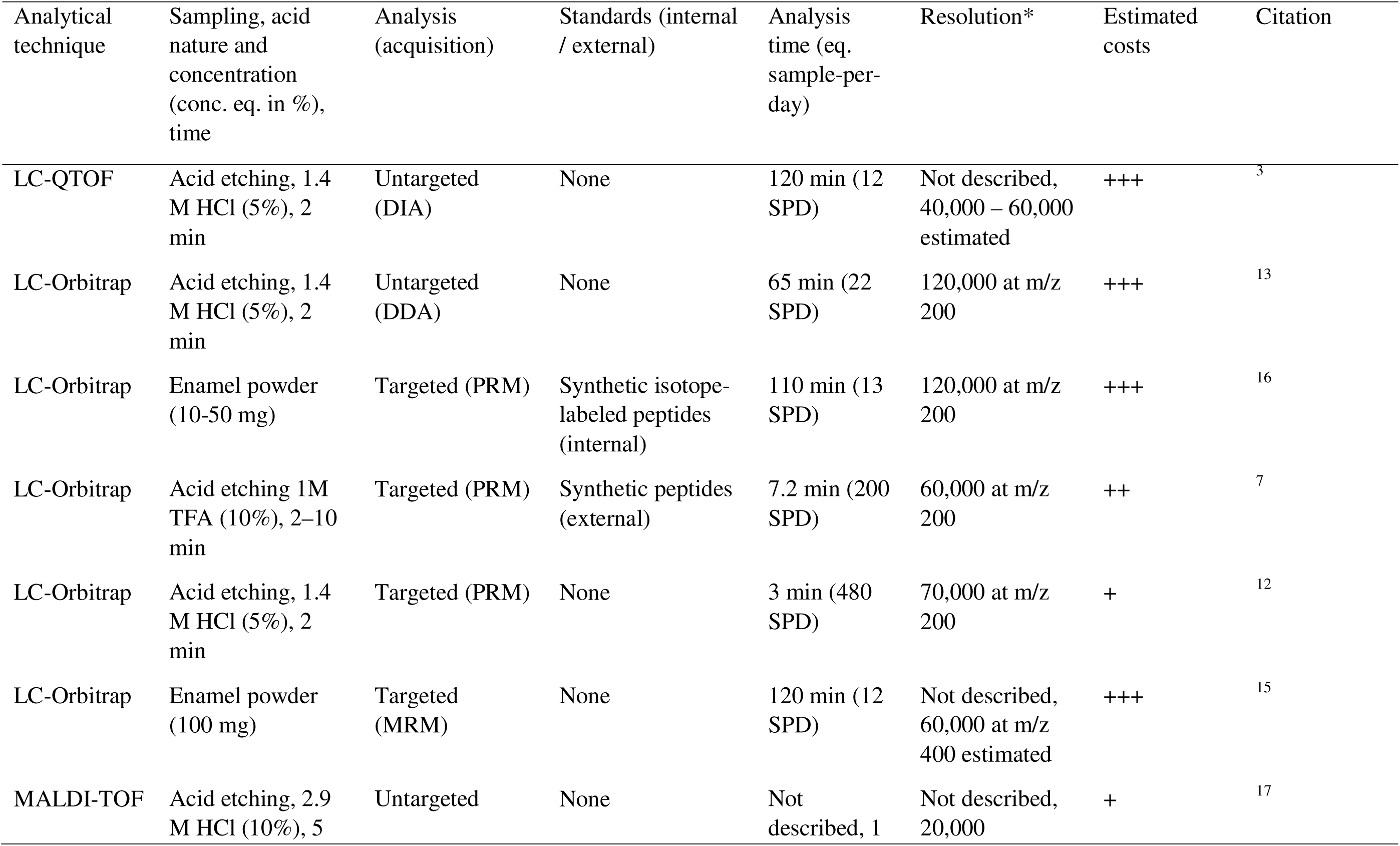
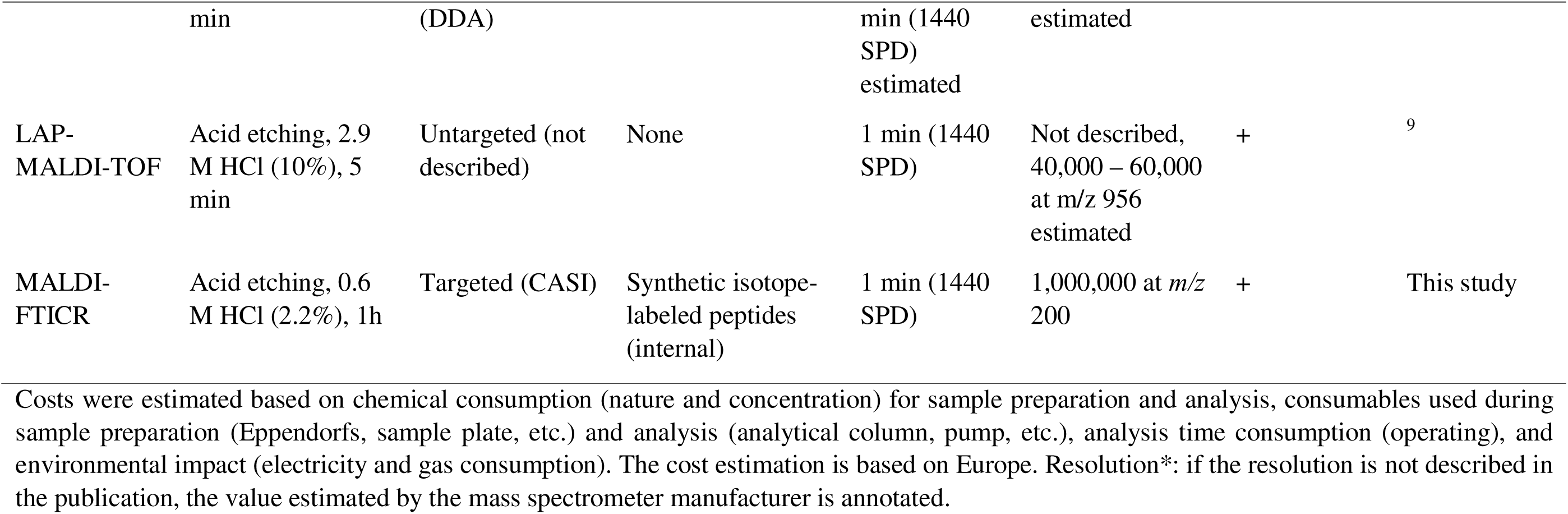
Comparison of currently available methods for biological sex estimation by proteomics.

The use of the CASI acquisition method with a Fourier transform ion cyclotron resonance (FTICR) mass spectrometer provides a low detection limit^26^. It has been shown that the CASI method yields results similar to those obtained using LC–HRMS for the analysis of archaeological bone samples^27^. We used two peptides, m/z 889.4491 (SM(ox)IR*PPY) for the AMELY isoform and m/z 1089.5621 (SIR*PPYPSY) for the AMELX isoform, as standards from the publication by Koenig *et al.*^7^. The heavy/light peptide ratio allows us to eliminate samples with too low an intensity for light or heavy peptides. The use of these peptides improves the confidence in biological sex estimation^7^. It has been shown that MALDI allows for extremely accurate ratio measurements, which are similar to those obtained by MRM analysis^28^. Adding a heavy peptide for the mass *m/z* 829.4590 would be useful for improving confidence in the results. The unique disadvantage of this method is the high cost of synthetic isotope-labeled peptides used for internal quantification (approximately €500 per peptide) compared to the synthetic peptides used for external quantification (approximately €150 per peptide).

It is important to note that the biological methods for estimating the sex of individuals has weaknesses. (i) It should be emphasized that female sex estimation carries the highest risk of false positives when data quality is low and AMELY signal intensity is weak. In 1992, Salido *et al.*^29^ have reported that transcription of amelogenin gene from the human Y chromosome may account for only approximately 10% of total amelogenin transcription. In 2019, Parker *et al.*^30^ demonstrated that a quantitative proteomic approach improves the estimation of false-positive rates and proposed a probabilistic method for sex estimation. However, the authors also highlighted the limitations of this approach, particularly due to the weak AMELY signal. More recently, in 2024, Koenig *et al.*^7^ developed a quantitative method based on LC-HRMS, incorporating a statistical framework specifically designed for the positive identification of female individuals. In this context, validation thresholds were applied in the present study to reduce false positives, relying on internal standards to strengthen the robustness of interpretations. (ii) Sex estimation by mass spectrometry also faces an additional limitation related to the possible deletion of the Y-chromosomal amelogenin gene. Such deletions have been reported in populations worldwide. Although this event remains rare—generally occurring in fewer than 1% of individuals and often well below 0.1%^31,32^—significantly higher frequencies have been documented in biological males from certain populations of the Indian subcontinent. In these cases, the absence of AMELY may lead to the misclassification of XY individuals as female when relying solely on proteomic detection. In addition, other biological conditions affecting the sex chromosome complement—such as sex chromosome trisomies, including Klinefelter syndrome (XXY)—may influence the results. These biological variations constitute important constraints that must be taken into account and highlight the need to interpret proteomics sex estimation within a broader genetic and anthropological context.

### Comparison of Morphological versus Proteomics Sex Estimation

With regard to the comparison of morphological sex assessment performed by different authors and biological sex estimation by proteomics analysis, the following observations can be made.

For adult individuals from the Mikulčice site, sex was assessed for all individuals in our sample (n = 30) by Stloukal in the 1960s^19,20^ using the methods available at that time^33–36^. Remarkably, despite the lack of sophisticated methods available today, only three out of the 30 cases (H392, H493, and H1082) were assigned a sex different from that estimated by proteomics analysis in the present study.

Regarding subsequent revisions, Zazvonilová *et al.*^21^ however, they were able to assess the sex of only two individuals from our sample. For the remaining individuals, the application of their selected methods^37–41^ was not possible. Their approach relied primarily on the pelvic bone, with the skull used preferentially in cases where the pelvis was not preserved.

Brůžek and Velemínský assessed sex using primary and secondary sex diagnosis^22^ for only a subset of individuals from Mikulčice, with 11 individuals overlapping with our sample. In one case, sex was not assessed, and the individual was marked with a question mark, whereas in another case (H1082), sex was assigned differently from that subsequently confirmed by proteomics analysis.

Notably, in one individual from our sample (H1082), both Stloukal^19,20^ and Brůžek with Velemínský—each highly experienced observers—confidently assigned the same sex, which was shown by proteomics analysis in the present study to be incorrect. Thus, although adult sex assessments made by experienced anthropologists were confirmed proteomically in the majority of cases, these findings demonstrate that proteomics approaches occupy an important and complementary role in anthropological research. This is particularly true in cases where the skeletal remains of adult individuals are poorly preserved, the pelvis is absent, and only fragmentary elements of other bones are available.

Proteomics sex estimation is particularly important in the case of non-adult individuals, for whom accurate and reliable sex assessment using morphological methods is not possible because the development of sexually dimorphic traits is incomplete. In this study, we analyzed 100 non-adult individuals from the Rajhrad site. Proteomics analysis identified 48 males and 51 females, while the biological sex could not be identified for one individual whose peptidic signal was too weak.

Although sex assessment is not routinely performed in non-adult individuals, sex was assessed for a total of 13 individuals in our sample by Pospíšilová^24^ using metric measurements of permanent teeth following the method proposed by Cardoso *et al.*^25^. Proteomics analysis confirmed the accuracy of these estimates in 11 of the 13 individuals, which can be considered a high success rate and indicates the good reliability of this method. Strong concordance between this metric approach and genetic sex estimation in non-adult individuals has also been demonstrated in a recent study^42^ and the method has been successfully applied in other contemporary research as well^43^.

## Conclusion

The newly developed high-throughput method for targeted paleoproteomics sex estimation using MALDI-CASI-FTICR mass spectrometry successfully estimated the biological sex of 129 to 130 adult and juvenile individuals from medieval Great Moravia. This approach combines the advantages of all methods commonly used to date, offering ultra-high resolution, substantially reduced processing times, applicability to large archaeological sample sets, and minimally invasive sampling. Internal quantification based on heavy/light peptide ratios further enhances the accuracy of biological sex estimation, ensuring reliability even in the presence of instrumental variations. The unique disadvantage of MALDI-CASI-FTICR is the high cost of synthetic isotope-labeled peptides used as the internal standard compared to the synthetic peptides used as the external standard in LC-PRM-HRMS.

The possibility of proteomics sex estimation in non-adult individuals opens up new opportunities for anthropologists and archaeologists, including more accurate reconstructions of past population structures, improved analyses of mortality patterns and sex-specific burial practices, and the investigation of social organization, kinship, and gender-related behavior in past societies, which have previously been inaccessible due to the inability to reliably determine sex in juvenile remains.

The method developed here still needs to be improved for the field of forensic medicine. The cohort should be expanded in order to refine specificity and false positive and negative rates. It is also important to have cohorts of individuals whose sex is known for different periods in order to determine the appropriate measures for each time period.

## Supporting Information

Supporting Information S1 (.docx file) contains Figure S1: MS spectra of modern male reference analyzed by MALDI-CASI-FTICR and MALDI-FTICR MS, Figure S2: Log_2_ ratio of intensities for the peptide *m/z* 889.4491 / *m/z* 879.4393 and *m/z* 1079.5520 / *m/z* 1089.5621 for 28 adult modern teeth, Figure S3: Log_2_ ratio of intensities for the peptide *m/z* 889.4491 / m*/z* 879.4393 and *m/z* 1079.5520 / *m/z* 1089.5621 for (A) Mikulčice site and (B) Rajhrad site.

Supporting Information S2 (.xlsx file) contains Table S1, S2: Log_2_ intensities of AMELX, Y, Ratio 889/879, Ratio 1089/1079 and sex estimation for both MALDI-FTICR MS and MALDI-CASI-FTICR-MS for Rajhrad and Mikulčice site respectively

## Ethics

This work did not require ethical approval from a human subject or animal welfare committee

## Declaration of AI use

Artificial intelligence was used for grammatical correction of the manuscript.

## Author contributions

Conceptualization: FB, APK, JB, MM; Data Curation: FB, MM; Formal Analysis: FB, LG, MM; Funding Acquisition: FB, CR, MM; Investigation: FB, APK, LG, MH, CR, PV, MM; Methodology: FB, LG, MC, JB, MM; Project Administration: MM; Resources: BB, KA, PV, CR, JB; Software: FB, LG, MH, MM; Supervision: MM; Validation: FB, APK, MM; Visualization: FB, LG, MM; Writing – Original Draft Preparation: FB, APK, MM; Writing – Review & Editing: FB, APK, LG, MH, BB, KA, CR, PV, MM

## Data availability

The mass spectrometry proteomics data have been deposited to the Zenodo Archive (https://zenodo.org/) via the Zonodo partner repository with the data set identifier 117817489.

## Acknowledgements

The authors are particularly grateful the financial support from the French national infrastructure INFRANALYTICS (FR2054 CNRS) for supporting the FTICR MS facility and the CNRS - Mission pour l’Interdisciplinarité for the funding of the EvoArPalMS project (2024-2025). This work was also supported by the Ministry of Culture of the Czech Republic (DKRVO 2024-2028/7.I.c, 00023272). The mass spectrometers were funded by the University of Lille, the CNRS, the Région Hauts-de-France and the European Regional Development Fund.

