## Supporting Information S1 for "High-throughput targeted paleoproteomics sex estimation on medieval Great Moravia individuals using MALDI-CASI-FTICR mass spectrometry"

^†^ Deceased

^*^ (MM)

**Figure S1**: MS spectra of modern male reference analyzed by MALDI-CASI-FTICR (up) and MALDI-FTICR MS (down). A) Full MS spectra and B) zoom in on the spectrum for the mass range m/z 815-1,115.	3

**Figure S2**: Log2 ratio of intensities for the peptide m/z 889.4491 / m/z 879.4393 and m/z 1079.5520 / m/z 1089.5621 for 28 adult modern teeth.	4

**Figure S3**: Log2 ratio of intensities for the peptide m/z 889.4491 / m/z 879.4393 and m/z 1079.5520 / m/z 1089.5621 for (A) Mikulčice site and (B) Rajhrad site.	4

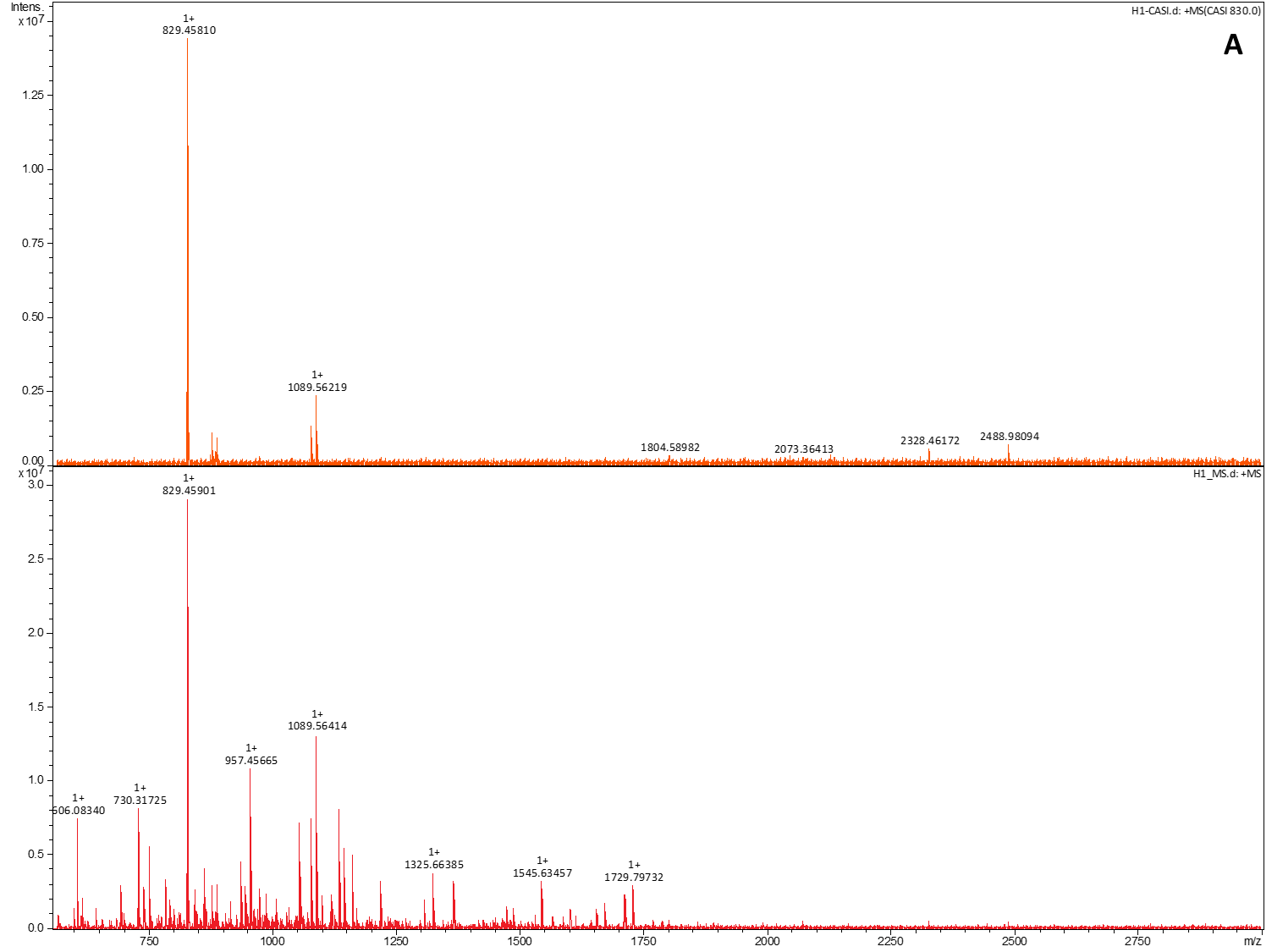

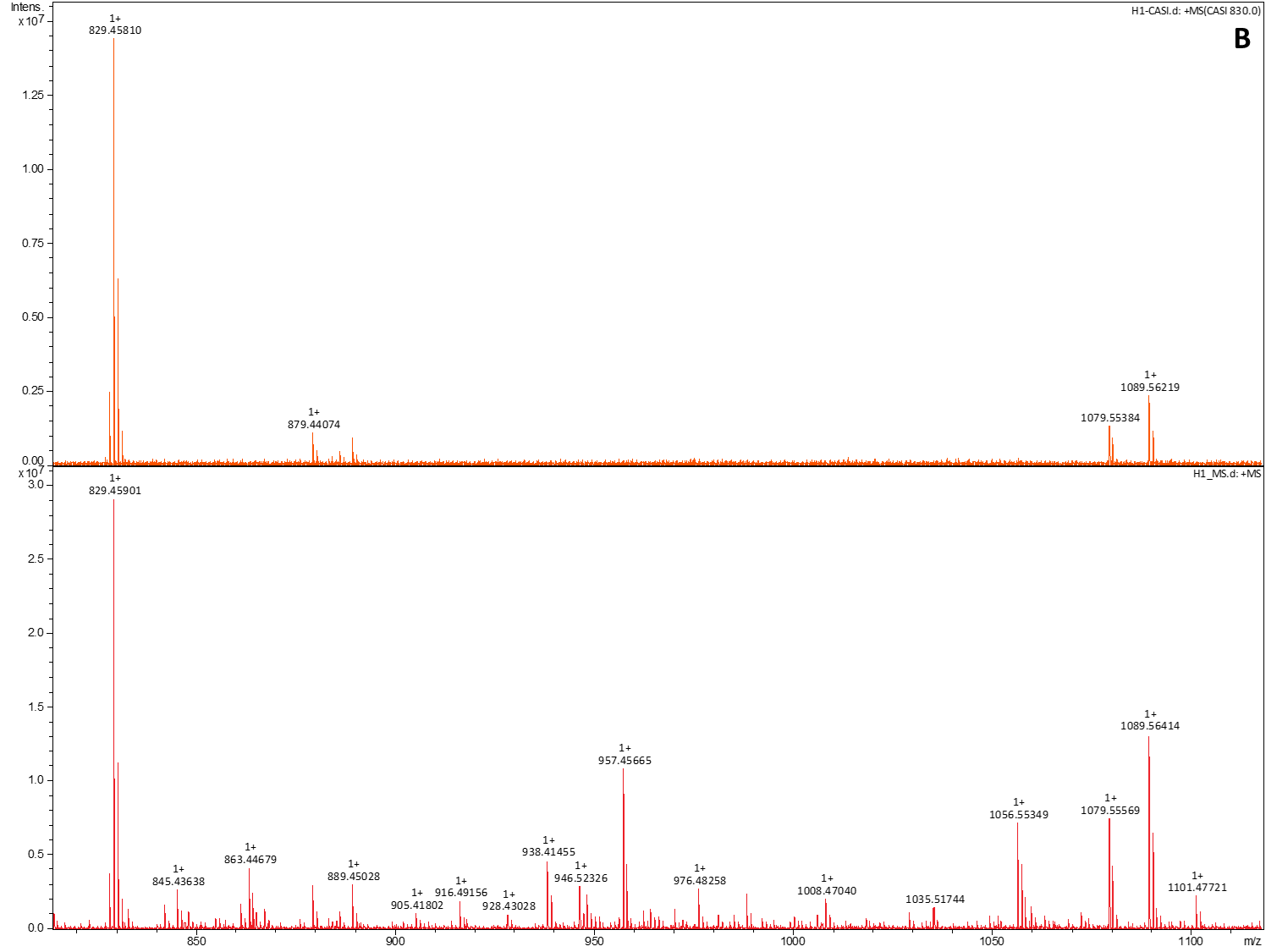

**Figure S1**: MS spectra of modern male reference analyzed by MALDI-CASI-FTICR (up) and MALDI-FTICR MS (down). A) Full MS spectra and B) zoom in on the spectrum for the mass range *m/z* 815-1,115.

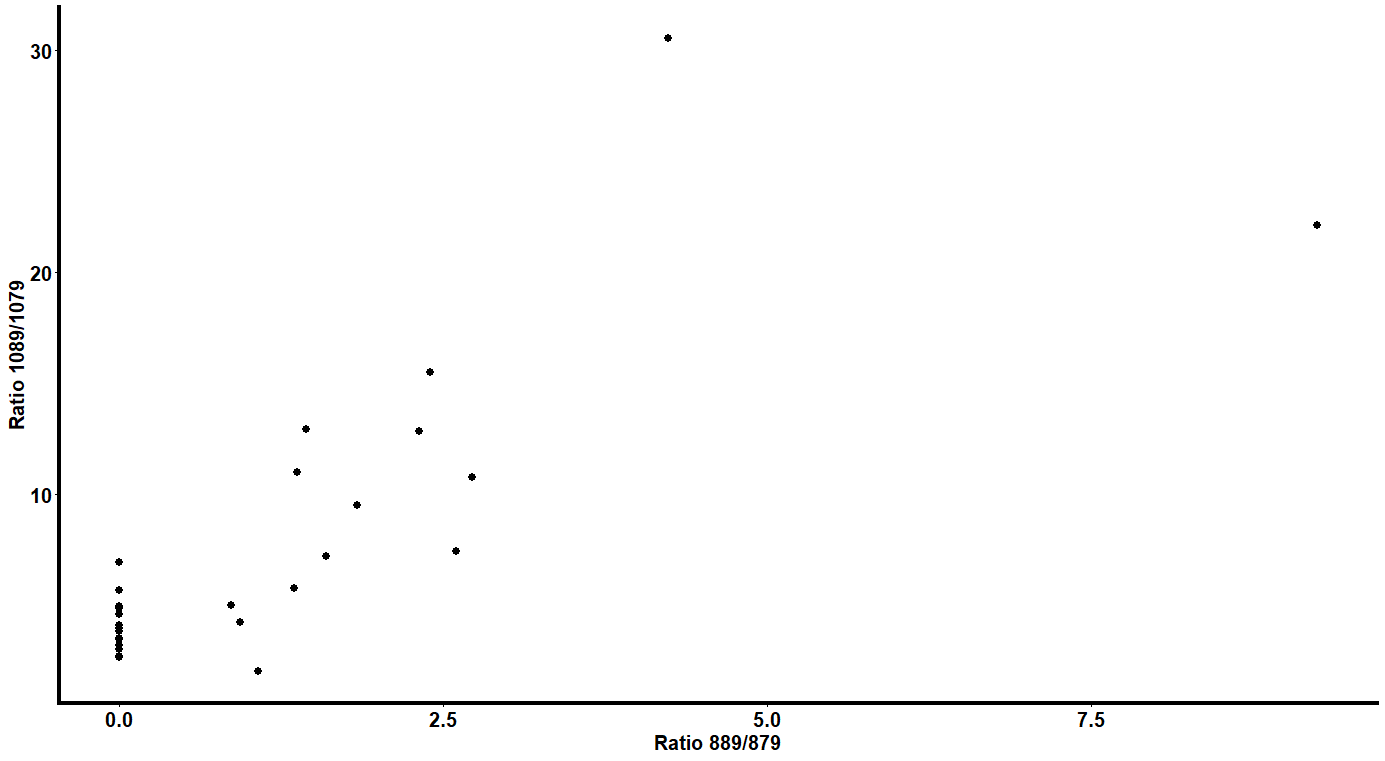

**Figure S2**: Log_2_ ratio of intensities for the peptide *m/z* 889.4491 / *m/z* 879.4393 and *m/z* 1079.5520 / *m/z* 1089.5621 for 28 adult modern teeth.

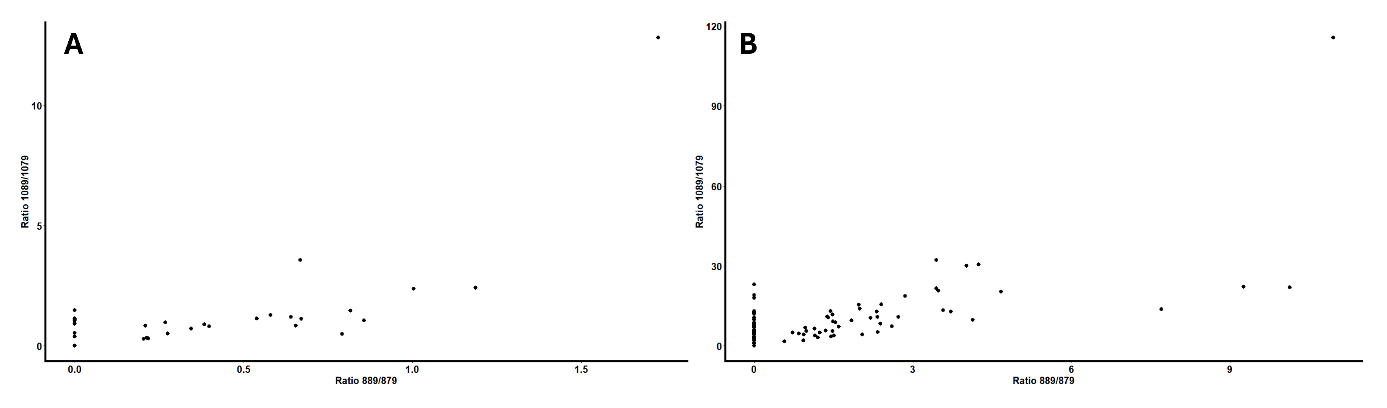

**Figure S3**: Log_2_ ratio of intensities for the peptide *m/z* 889.4491 / *m/z* 879.4393 and *m/z* 1079.5520 / *m/z* 1089.5621 for (A) Mikulčice site and (B) Rajhrad site.
